# TPCAV: Interpreting deep learning genomics models via concept attribution

**DOI:** 10.64898/2026.01.20.700723

**Authors:** Jianyu Yang, Shaun Mahony

## Abstract

Interpreting genomics deep learning models remains challenging. Existing feature attribution methods are largely restricted to one-hot DNA inputs and therefore cannot assess the influence of more general genomic features such as chromatin states or genomic repeats. Concept attribution methods offer an input-agnostic global interpretation framework, yet they have not been systematically applied to interpret neural network applications in genomics.

We present the first application of concept attribution to interpret genomics deep learning models by adapting the Testing with Concept Activation Vectors (TCAV) method. We improve upon the original TCAV method by incorporating a PCA-based decorrelation transformation to address correlated and redundant embedding features commonly observed in genomics deep learning models, resulting in the Testing with PCA-projected Concept Activation Vectors (TPCAV) approach. We also introduce a strategy for extracting concept-specific input attribution maps. We evaluate our approach by interpreting influential biological concepts across a diverse set of genomics models spanning multiple input representations and prediction tasks.

We demonstrate that TPCAV provides comparable motif feature interpretation to TF-MoDISco on one-hot encoded DNA-based transcription factor binding prediction models. TPCAV also enables robust interpretive analysis of how more general biological concepts such as repetitive elements and chromatin state annotations contribute towards predictions. TPCAV uniquely generalizes to interpret features learned by tokenized foundation models as well as models incorporating chromatin signals as inputs. We further show that TPCAV can identify representative regions associated with specific concepts, motivating downstream investigation of distinct regulatory mechanisms. TPCAV provides a flexible and robust complement to existing model interpretation techniques.

## Introduction

Deep learning has become a central tool in genomics, achieving state-of-the-art performance across a wide range of predictive tasks. A prominent class of these models – sequence-to-function models – typically take one-hot-encoded DNA sequences as input to predict diverse genomic features, including transcription factor (TF) binding^1,2^, regulatory signal tracks^3,4^, and RNA expression levels^5^. In recent years, both the scale and accuracy of sequence-to-function models have advanced dramatically. Notable examples include BPNet^1^, SpliceAI^6^, Borzoi^5^, and Enformer^3^, with the recent AlphaGenome model capable of predicting hundreds of genomic tracks across a 1-Mbp input window at single base pair resolution^4^. In parallel, efforts to incorporate additional input modalities, such as chromatin accessibility and histone modifications, have demonstrated further performance gains and provided deeper biological insight^7,8^.

Despite these advances, our understanding of what deep learning models learn, and how to translate that knowledge into meaningful biological insight, remains limited. A major challenge lies in the lack of unified approaches for interpreting the features learned by increasingly sophisticated architectures. Deep learning model interpretation in genomics has traditionally relied on feature attribution techniques, which fall into two broad categories: local and global attribution. Local methods, such as DeepLIFT^9^, provide base-level attribution scores for individual inputs and are widely used, but they primarily explain context-specific model behavior at individual genomic loci and are therefore difficult to summarize. In contrast, global feature attribution methods aim to capture the broader, high-level patterns learned by a model across many examples, offering a more comprehensive view of model behavior.

In genomics, TF-MoDISco^10^ is currently the most widely used global interpretation method for models operating on one-hot encoded DNA inputs. TF-MoDISco extracts informative sequence motifs from attribution scores and has demonstrated robust performance across multiple studies, providing valuable insights into model behavior. However, its reliance on one-hot encoding limits its applicability, preventing interpretation of models that use alternative input formats such as tokenized DNA. Moreover, for other input modalities, such as genomic signal tracks, there are currently no effective global interpretation methods capable of identifying the patterns a model relies on and summarizing them into testable biological hypotheses. With only local feature attribution available, interpretation is restricted to predefined genomic regions, offering only a narrow view of model decision-making. This significantly limits our ability to fully leverage high-performing deep learning models across diverse genomic tasks.

Concept-based attribution methods, such as Testing with Concept Activation Vectors (TCAV)^11^, provide an input-agnostic approach to interpreting deep learning models. Rather than summarizing local attribution scores, concept-based methods use sets of intentionally selected examples to represent high-level concepts that can be used to query trained deep learning models. In genomics, decades of research have established rich prior knowledge about the regulatory mechanisms underlying gene expression, potentially making concept-based attribution well suited for integrating known biological concepts into model interpretation. However, concept attribution methods have mainly been developed in the context of image classification tasks and have not yet been applied to interpret genomics deep learning models.

In this study, we adapt TCAV for genomics applications and introduce TPCAV (Testing with PCA-projected Concept Activation Vectors), which incorporates a PCA transformation into the TCAV workflow to address challenges associated with redundant, correlated features in the embedding space. We also introduce several additional features to adapt concept attribution to genomics deep learning model interpretation, including a comprehensive curated genomics concept database, a new metric to rank motif concept importance for a given task, and a concept-specific input attribution dissection strategy that validates concepts and ranks input genomic regions according to their use of a given concept. By applying TPCAV to various genomics deep learning models and tasks, we demonstrate that TPCAV offers intuitive and comprehensive interpretation of the importance of known biological concepts. TPCAV is highly generalizable across neural network architectures and input modalities, enabling interpretation of tokenized language models and other settings for which no effective global interpretation techniques currently exist.

## Results

### Testing with PCA-projected Concept Activation Vectors (TPCAV)

The TCAV concept-based attribution method investigates the sensitivity of deep learning model predictions to human-defined concepts^11^. Each concept is defined by a set of selected examples that each prominently contain features of the corresponding concept. For example, in the original application of TCAV to image classification models, a concept such as “stripiness” would be defined via a collection of images that each prominently contain examples of striped patterns^11^. In our application to genomics, we define DNA motif concepts using collections of sequences containing embedded motif instances. Concepts can also be defined using collections of genomic sequences that overlap particular genomic annotations (e.g., promoters or repeat elements) or using regions that display particular cell-specific regulatory activities (e.g., chromatin accessibility or chromatin states).

In TCAV, sets of concept examples and random (non-concept) examples are presented to the trained model, and a linear classifier is trained to distinguish between them based on vectors of node activations at a selected intermediate layer. The investigated layer is typically chosen to be close to the final layer as concepts usually represent high-level features that are learnt relatively later in the model. The vector orthogonal to the decision hyperplane of the classifier then represents the direction of the concept in the layer’s embedding space and is termed the “Concept Activation Vector” (CAV). The classification performance (measured via F-score) of this linear classifier is one indication of whether the neural network has embedded features relating to the concept. By multiplying the CAV by a given data point of interest’s attribution scoring vector from the same model layer, we can assess the sensitivity of model predictions to changes in the input in the direction of the CAV, conditioned on the test regions. Concept influence in a model can thereby be assessed across an entire class of inputs by calculating the fractions of data points for which the CAV increases or decreases model output scores, termed the TCAV score. Note that TCAV does not require any modification to the model architecture or training procedure and so it can be applied to any trained model for which we can investigate node activities at an intermediate layer.

To investigate whether TCAV can be applied to interpret genomics models, we first developed a TF binding prediction model, consisting of a convolutional tower followed by transformer encoder layers and a final dense prediction layer (see **Methods** for details). The task is to predict whether a 256-bp region is bound by a TF, using a 1,024-bp one-hot encoded DNA sequence centered on the test region as input. We downloaded the ENCODE Dream Challenge datasets^12^ and trained a separate model for each TF in the HepG2 and H1-hESC cell lines. Most models achieved strong performance; models with a test-set auPRC below 0.8 were excluded from downstream analyses (**Supplementary Table S1**).

We next investigated the sensitivity of each trained TF binding prediction model to the relevant cognate TF-DNA motif concepts. Specifically, we created a library of DNA motif concepts by embedding instances of known human TF binding motifs into randomly sampled sequences from the human genome, taking the non-redundant clustered motif collection from the Vierstra lab as our source of known motifs^13^ (see **Methods**). The CAVs were calculated using activations from the neural network layer immediately preceding the final dense layer in each model. In most cases, TCAV found the expected cognate motif concept to be a positive influence in the corresponding TF binding models (**Figure 1b**), suggesting that TCAV identifies informative concepts. However, we noted some unintuitive exceptions. For example, TCAV finds that the CTCF motif concept contributes negatively to CTCF binding predictions in HepG2 cells (**Figure 1b**). Since CTCF binding is highly positively associated with the presence of the CTCF cognate binding motif in all cell types, it appears that TCAV has misidentified the sign of the cognate motif concept influence in this case. A similar issue is also observed in the NANOG prediction model in H1-hESC cells, where TCAV finds NANOG’s cognate motif concept as a negative contributor (**Figure 1b**).

**Figure 1:**
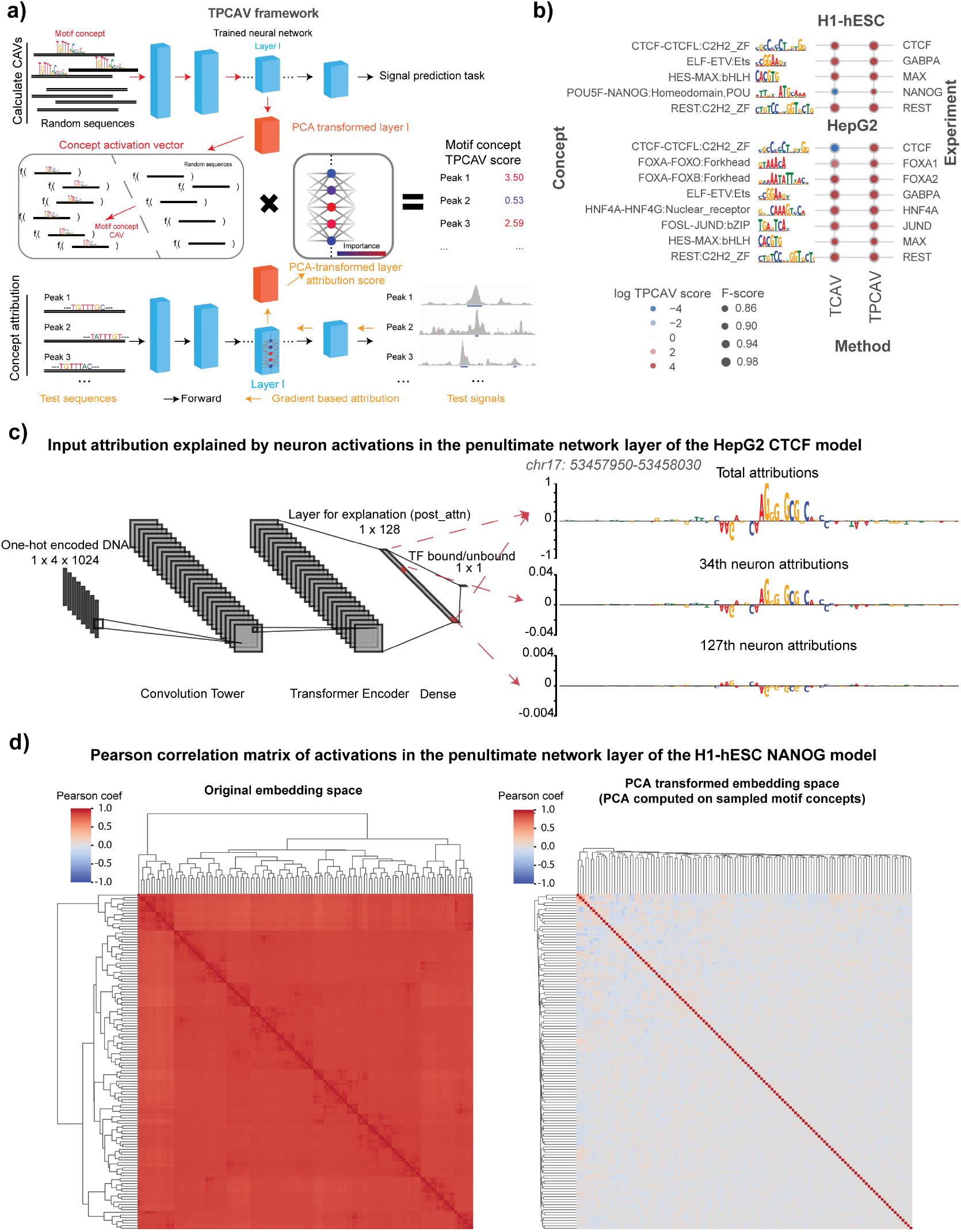
Overview of the TPCAV algorithm and comparison with TCAV. **a)** TPCAV adds an additional PCA transformation to the feature embedding space, which decorrelates the feature dimensions. **b)** Comparison of TCAV and TPCAV when determining the influence of cognate motif concepts in sequence-specific TF binding prediction models trained on H1-hESC and HepG2 cell line datasets from the ENCODE DREAM Challenge. **c)** Input DeepLIFT attribution scores from the HepG2 CTCF prediction model at a CTCF binding site (chr17: 53457950-53458030). Top panel: combining all activations in the penultimate neural network layer; middle panel: using attributions associated only with the 34^th^ neuron activations; bottom panel: using attributions associated only with the 127^th^ neuron activations. **d)** Pearson correlation of neuron activations from the penultimate layer in NANOG peaks and random regions before (left) and after (right) PCA transformation.

Examining the properties of these two binding prediction models in further detail, we determined that TCAV’s unintuitive behavior is likely due to the presence of multiple correlated embedding space features in the model that redundantly encode the cognate motif. For example, in the HepG2 CTCF prediction model, we found that the chosen embedding layer contains multiple nodes that respond to the presence of the CTCF cognate motif, but some respond positively while some respond negatively (**Figure 1c**). Overall, the summation of all node activations results in CTCF motif instances positively contributing to the prediction, so the presence of redundant, contradictory nodes is not necessarily an issue for the performance of the TF binding model. However, since the redundant node activities are all correlated, TCAV’s linear classifier will sometimes use negatively-contributing node activities to classify concept examples, and therefore the resulting CAV will point in the opposite orientation to its expected direction.

To resolve TCAV’s issues with highly redundant and correlated embedding space features, we incorporated an additional PCA transformation into the TCAV algorithm, yielding the Testing with PCA-projected Concept Activation Vectors (TPCAV) approach (**Figure 1a**). Since PCA transformation is linear and reversible, it can decorrelate the embedding space and keep the original forward pass intact. The PCA transformation is applied to sampled examples for each candidate concept. By visualizing the correlations between node activations across TF peaks and random genomic regions demonstrates that this transformation effectively decorrelates the embedding space. (**Figure 1d, Supplementary Figure S1a**). Applying TPCAV to interpret the influence of the same cognate motif concepts on the trained TF binding models shows that it correctly identifies all cognate motif concepts as positive influences, improving the reliability of the original method (**Figure 1b**).

### TPCAV provides comprehensive interpretation of DNA features

When we applied TCPAV to assess the influence of a wider collection of human TF-DNA binding motifs on the trained TF binding models, we found that substantial numbers of motif concepts are classifiable via TPCAV with high F-scores (**Supplementary Table S2**). To better assess the relative importance of these motif concepts, we introduce a new metric – the motif concept sensitivity score – defined as the corrected area under the curve (AUC) of each motif concept classifier’s F-score across varying numbers of motif insertions within concept examples. The correction is performed using linear regression to regress out the effect of GC information content, thereby accounting for bias introduced by motif GC content (see **Methods, Figure 2a**). This metric enables us to rank motif concepts according to how strongly a model responds to increasing numbers of motif instances. In most TF binding prediction models, the corresponding cognate motif ranks among the top 3 concepts (**Figure 2b**). However, in some cases, the expected cognate motif ranks lower than other DNA-binding motifs. For example, the MAX-related motif concept USF-BHLBH:bHLH ranks 6^th^ in the HepG2 MAX prediction model (**Table S2**). In this and other similar cases, the top-ranked motif concepts often correspond to either cell line-specific TFs (e.g., the HNF4A cognate motif in HepG2), or GC-rich motifs (e.g., ZNF-PAX:C2H2_ZF), suggesting that the models may partially rely on cell-specific motifs or GC-content as general predictive features of regulatory regions in addition to the cognate motifs (**Table S2**). Furthermore, cognate motif concepts exhibit strong experiment specificity; they tend to rank highly only in the ChIP-seq experiment corresponding to the appropriate TF (**Figure 2b, Table S2**). We also examined motif concept CAVs to investigate how they relate to each other in the embedding space. Using the HepG2 FOXA1 model as an example, we computed the cosine similarity matrix among the top motif-concept CAVs and performed hierarchical clustering. As expected, motif concepts with similar position weight matrices clustered together, indicating that TPCAV reliably captures coherent concept directions for related sequence motifs (**Supplementary Figure S1b**).

**Figure 2:**
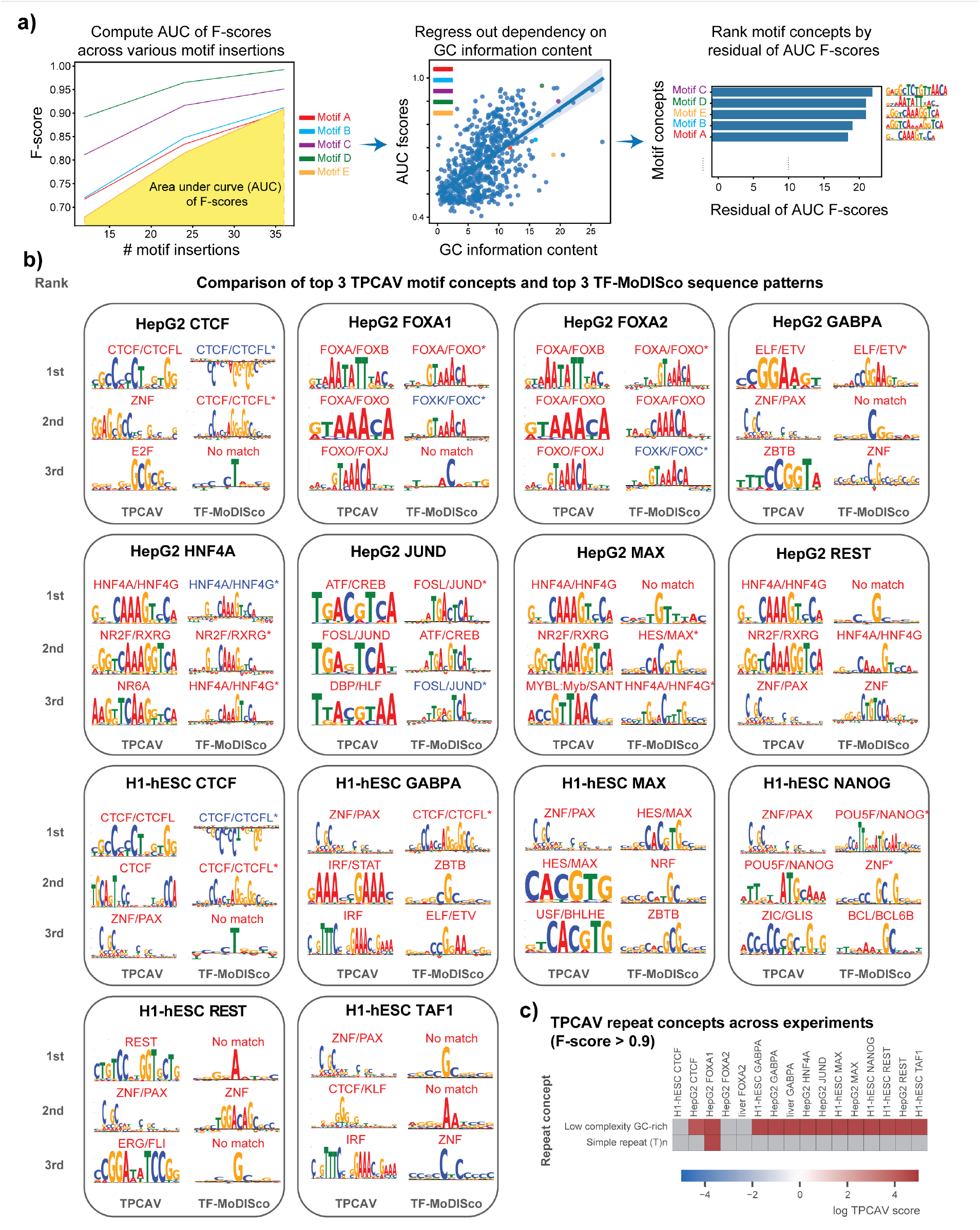
TPCAV offers comprehensive interpretation results for DNA motifs and repeats. **a)** Illustration of motif concept sensitivity score computation by first obtaining motif concept AUC of F-scores across various numbers of motif insertions, then correcting the effect of motif GC information content. **b)** Comparison of the top three identified motif’s from TPCAV (ranked by motif concept sensitivity score) and TF-MoDISco (ranked by number of seqlets) on models trained in HepG2 and H1-hESC. Significant matches to database motif are marked by “*” for TF-MoDISco. Motifs with blue names appeared in TF-MoDISco’s negative motif set. **c)** DNA repeat concepts with F-score > 0.90 across models trained in HepG2 and H1-hESC.

To further validate the interpretation results obtained from TPCAV, we performed TF-MoDISco analysis on each model. The sequence patterns identified by TF-MoDISco were largely consistent with the cognate motifs of the corresponding TFs, with most cognate motifs appearing among the top 3 contributing sequence patterns (**Figure 2b**). It is notable that some TF-MoDISco motifs visually corresponding to cognate motifs are not deemed significant matches by Tomtom^14^, which could be explained by noisy flanking sequences.

TPCAV yields uniformly positive influence scores for the top motif concepts (**Figure 2b**). In contrast, TF-MoDISco, while typically recovering cognate motifs among the strongest positive patterns, often also identifies the same cognate motif as a top negative pattern. For example, the CTCF cognate motif appears as both the highest positive and highest negative pattern in the CTCF models for H1-hESC and HepG2. Similar patterns are observed for FOXA1 and FOXA2 in HepG2 and for JUND in HepG2 (**Figure 2b** and **Supplementary Figure S2**). These discrepancies likely arise because the predictive models can sometimes assign negative attribution scores to certain redundant occurrences of the cognate motif or to sequences flanking the cognate motif. TF-MoDISco is designed to separately cluster positively and negatively contributing seqlets and therefore the method cannot provide a definitive global interpretation when the same sequence pattern appears in both groups. In contrast, TPCAV evaluates each concept globally and produces a single aggregated score, thereby avoiding this ambiguity. In addition, many of TF-MoDISco’s top sequence patterns include only a single highly contributing nucleotide, which may reflect the model’s use of local GC-content signals (**Figure 2b** and **Supplementary Figure S2**).

Because our TF binding prediction models use a relatively large 1,024-bp window to predict binding within the central 256-bp, we asked whether the models exhibit positional preferences for motif occurrences across the input sequence. We reconstructed each motif concept multiple times, each version placing four motif instances within a specific 128-bp subregion of the input window. We then examined the F-scores of these location-specific motif concepts. The results show that even the top motif concepts are classifiable only when inserted around the central 256-bp subregion (**Supplementary Figure S3**). This indicates that the models indeed display strong positional preferences for motif concepts and that TPCAV can effectively capture these positional biases.

Beyond DNA motif-related concepts, TPCAV can be applied to assess the influence of annotation-based concepts that do not necessarily have uniform or consistently definable sequence characteristics. For example, repeat elements constitute a substantial portion of mammalian genomes and represent an important category of DNA features. To assess whether any repeat families contribute to predictions in the trained TF binding models, we applied TPCAV to evaluate repeat concepts defined via instances of each repeat class in the RepeatMasker database^15,16^. Across models, the most frequently detected repeat concept was the low-complexity GC-rich class (**Figure 2c, Supplementary Table S3**). Because GC-rich regions often correspond to CpG islands and are strongly associated with open chromatin, this suggests that the models may use GC-richness as a proxy for chromatin accessibility. The only other repeat concept that was consistently classifiable was the simple repeat (T)_n_, which showed a positive influence in the FOXA1 model trained on HepG2. FOXA1 and other Forkhead TFs have been shown to preferentially interact with T-rich simple repeats^17,18^.

In summary, TPCAV reliably identifies cognate motif concepts as positively influential in TF binding prediction models, and its results are generally consistent with those obtained from TF-MoDISco. While TF-MoDISco excels at discovering motif patterns *de novo*, TPCAV provides a complementary direct interpretation of known motif concepts. By examining the cosine similarities between CAVs, TPCAV enables assessment of relationships and correlations among motif concepts within the model. Moreover, the flexibility of TPCAV allows us to test the positional preference of motif concepts and the influence of annotation-based concepts that are not necessarily defined by a single sequence feature.

### TPCAV is an input format agnostic global feature attribution approach

Although TF-MoDISco can discover important motif features *de novo*, its applicability is limited to models operating on one-hot-encoded DNA inputs. In contrast, concept-based methods require only example sets that represent each concept, without the need to define the concept mathematically or constrain the input representation. This flexibility allows TPCAV to evaluate a wide range of concept types and to generalize across diverse model architectures.

Large language models have become increasingly popular for deep learning applications in genomics. Several studies have demonstrated that leveraging pre-trained DNA foundation models can substantially improve prediction performance, particularly when training data are limited^19–21^. These foundation models typically operate on tokenized DNA inputs, which reduce the number of model parameters and improve training efficiency. However, tokenization, while enabling large input windows and scalable model training, is incompatible with the one-hot input requirement of TF-MoDISco, preventing global interpretation of such models.

To address this gap, we first applied TPCAV to interpret a widely used DNA foundation model, DNABERT-2^19^. We finetuned the pre-trained DNABERT-2 foundation model to predict FOXA1 ChIP-seq peaks from the A549 cell line using random genomic regions as negatives. After finetuning, the DNABERT-2 model achieves an F-score of 0.86 and auPRC of 0.82 (baseline 0.51) on the evaluation dataset. For TPCAV interpretation, we used the embeddings of the pooler layer following the BERT encoder. TCPAV analysis of DNA motif concepts show that both the FOXA1 cognate motif and the JUND motif rank as top concepts and exert strong positive influence on FOXA1 peaks (**Figure 3a, Table S2**). These results are consistent with prior studies showing a strong dependence between FOXA1 and AP-1 (for which the JUND motif is representative) in A549 cells^22^. Our findings suggest that TPCAV can be effectively applied to interpret tokenized DNA foundation models.

**Figure 3:**
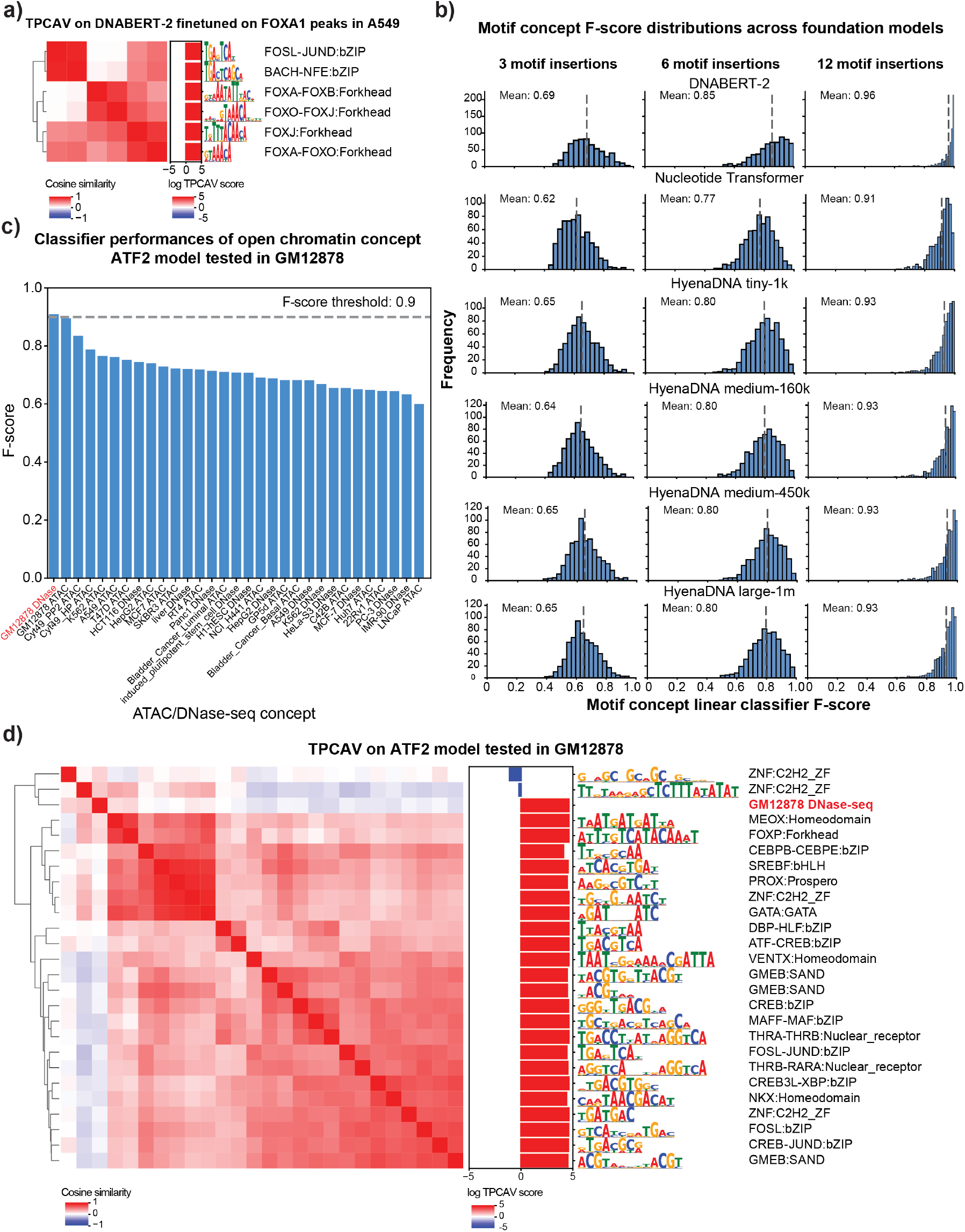
TPCAV is a generalizable feature attribution method. **a)** CAV cosine similarity heatmap and TPCAV scores of motif concepts (amongst top 30 motifs, with F-score > 0.9) in a DNABERT-2 model that is finetuned on FOXA1 peaks in A549 cells. **b)** Distributions of F-scores for motif concepts under 3, 6, 12 insertions in the embedding space of each foundation model. **c)** F-scores of open chromatin concepts in an ATF2 prediction model applied in GM12878 cell line. **d)** CAV cosine similarity heatmap and TPCAV scores of motif concepts (amongst top 30 with F-score > 0.9) and GM12878 DNase-seq peak concept (F-score > 0.9) in ATF2 prediction model applied in GM12878 cell line.

In addition to DNABERT-2, there are several types of foundation models available from different groups, including Nucleotide Transformer^23^ and HyenaDNA^24^. Although these pre-trained models offer advantages such as improved fine-tuning performance on small datasets and faster training, there is still no consensus regarding the optimal architecture or training strategy. A major barrier to gaining such insights is the lack of suitable interpretation methods for these models.

TPCAV interpretation and our biological concept database provides a practical framework for benchmarking foundation models. Useful DNA foundation models should learn a broad range of regulatory motif features. Therefore, TPCAV should be able to use the embedding space provided by higher levels of a foundation model to effectively classify a broad range of motif concepts from random sequences. To investigate how successfully foundation models encapsulate motif concepts, we trained TCPAV motif concept classifiers using 3, 6, and 12 motif insertions within a 512-bp input window and evaluated them in the embedding spaces of several pretrained foundation models. The resulting F-score distributions reveal that DNABERT-2 consistently outperforms the Nucleotide Transformer and HyenaDNA models by accurately distinguishing motif concepts across all insertion levels (**Figure 3b**). Comparison across HyenaDNA models with different parameter sizes revealed only minor differences in classification performance, suggesting that parameter count alone does not explain the model’s reduced ability to distinguish motif concepts (**Figure 3b**). To further characterize the relatively poorly classified motif concepts in each model, we clustered them using STAMP^25,26^. The concepts missed by DNABERT-2 tended to feature short, low-information motifs (**Supplementary Figure S4**), which naturally exhibit limited distinction from the control concept. In contrast, the Nucleotide Transformer and HyenaDNA models missed a relatively broader range of motif concepts, potentially reflecting differences in dataset composition or training strategies (**Supplementary Figure S4**). We further examined the top 30 motif concepts in each foundation model to assess whether individual models exhibit preferences for specific subsets of TF motifs. The results indicate that all models show a consistent preference for AT-rich motif concepts (**Supplementary Figure S5**).

These results demonstrate that TPCAV is an input-agnostic framework that can be applied to interpret foundation models operating on tokenized DNA sequence inputs. By evaluating the motif concepts across foundation models, we found that DNABERT-2 encodes regulatory motif information more effectively than Nucleotide Transformer and HyenaDNA.

### TPCAV can interpret input modalities beyond DNA sequence

Beyond sequence-only models, many groups have begun incorporating diverse genomic modalities, including ATAC-seq and histone modification ChIP-seq, to improve predictive performance or interpretive capacity^7,27,28^. To evaluate whether TPCAV can interpret multimodal architectures, we applied it to maxATAC, a TF prediction framework that integrates DNA sequence with ATAC-seq signal to enable cross-cell-type predictions^7^. In addition to the motif and repeat concepts used in our sequence-only models, we introduce open-chromatin concepts derived from DNase-seq and ATAC-seq peaks across multiple cell lines. To ensure that these concepts capture the effect of the chromatin accessibility signal rather than DNA sequence features associated with chromatin accessibility, each open-chromatin concept is constructed by pairing the DNase/ATAC-seq signal from a peak with a random DNA sequence.

We examined two maxATAC models, an ATF2 model tested in GM12878 and a RUNX1 model tested in K562. We choose the activations from the last max pooling layer for TPCAV interpretation. Across both models, the open-chromatin concept corresponding to the matched cell line was the most classifiable, as indicated by the highest concept-classifier F-scores (**Figure 3c, Supplementary Figure S6a**). We then investigated all classifiable concepts and visualized their relationships using cosine similarity heatmaps, resulting in three broad clusters of concepts (**Figure 3d**). The largest cluster consisted of motifs similar to the ATF2 cognate motif. The GM12878 DNase-seq concept was clearly separated from the motif cluster, indicating that the model treats chromatin accessibility signals as a distinct feature from sequence motifs. In addition, the GM12878 DNase-seq concept is closely clustered with the GC-rich concept ZNF:C2H2_ZF, which shows a negative influence on the model predictions (**Figure 3d**). Notably, this pattern contrasts with our observations in sequence-only TF binding models, in which GC-rich concepts generally show positive influences. This suggests that GC-richness may serve as a proxy for chromatin accessibility when only DNA sequence is available; however, once explicit chromatin accessibility information is available, GC-rich features are no longer beneficial for prediction.

Similarly, applying TPCAV to the RUNX1 binding prediction model in K562 revealed that the RUNX1 cognate motif concept, along with several related motif concepts, showed strong positive influence on model predictions. As in the ATF2 model, multiple GC-rich concepts contributed negatively. In addition, the K562 open-chromatin concept showed a clear positive influence, forming a direction distinct from the motif-based concepts (**Supplementary Figure S6b**).

In summary, TPCAV can readily evaluate a wide range of genomics concepts beyond DNA sequence inputs. Our results demonstrate that TPCAV is a highly generalizable and flexible method for global feature attribution in multimodal models.

### TPCAV applied to a multi-headed model reveals differential usage of concepts in the shared embedding space

Thus far, we have applied TPCAV to models that predict the binding probability of a single TF. Several deep learning architectures simultaneously predict multiple genomic signals, resulting in a substantially more complex embedding space. To evaluate how TPCAV performs in this setting, we applied it to BPNet, a base-pair resolution TF binding prediction model trained to predict four TFs – Oct4, Sox2, Klf4, and Nanog (OSKN) – in mouse embryonic stem (mES) cells^1^. BPNet includes eight prediction heads, corresponding to the read-count and probability profiles for each factor. For TPCAV interpretation, we used the activations from the bottleneck layer immediately preceding the branching prediction heads.

Consistent with the original study, several motif concepts relevant to OSKN factors are among the top-ranked concepts, including KLF-SP:C2H2_ZF, SOX:Sox, and POU5F-NANOG:Homeodomain (**Figure 4a, Supplementary Table S2**). In addition, relevant motif concepts are classifiable and clustered around the cognate motif concepts (**Figure 4b**). The activity patterns of the cognate motif concepts aligned with expectations, where SOX-related motif concepts are positive influences in Sox2 and Nanog count tasks, POU domain-related motif concepts are active in Oct4, Nanog, and Sox2 count tasks, and Klf4-related motif concepts are most active in the Klf4 count task. In addition, several highly repetitive motif concepts emerged, potentially reflecting varying GC-content preferences (**Figure 4c**). For profile prediction tasks, the overall concept activity patterns are similar to those observed in count prediction tasks. In general, a larger number of motif concepts positively contribute to profile prediction tasks than count prediction tasks. For example, in Sox2 prediction, many motifs clustered around the Sox2 cognate motif show positive contributions in the profile task, whereas the count task is influenced more specifically by the Sox2 cognate motif itself. This may reflect the inherently noisier nature of profile predictions, which likely require the model to attend to a wider range of sequence features similar to cognate motifs. (**Figure 4c**).

**Figure 4:**
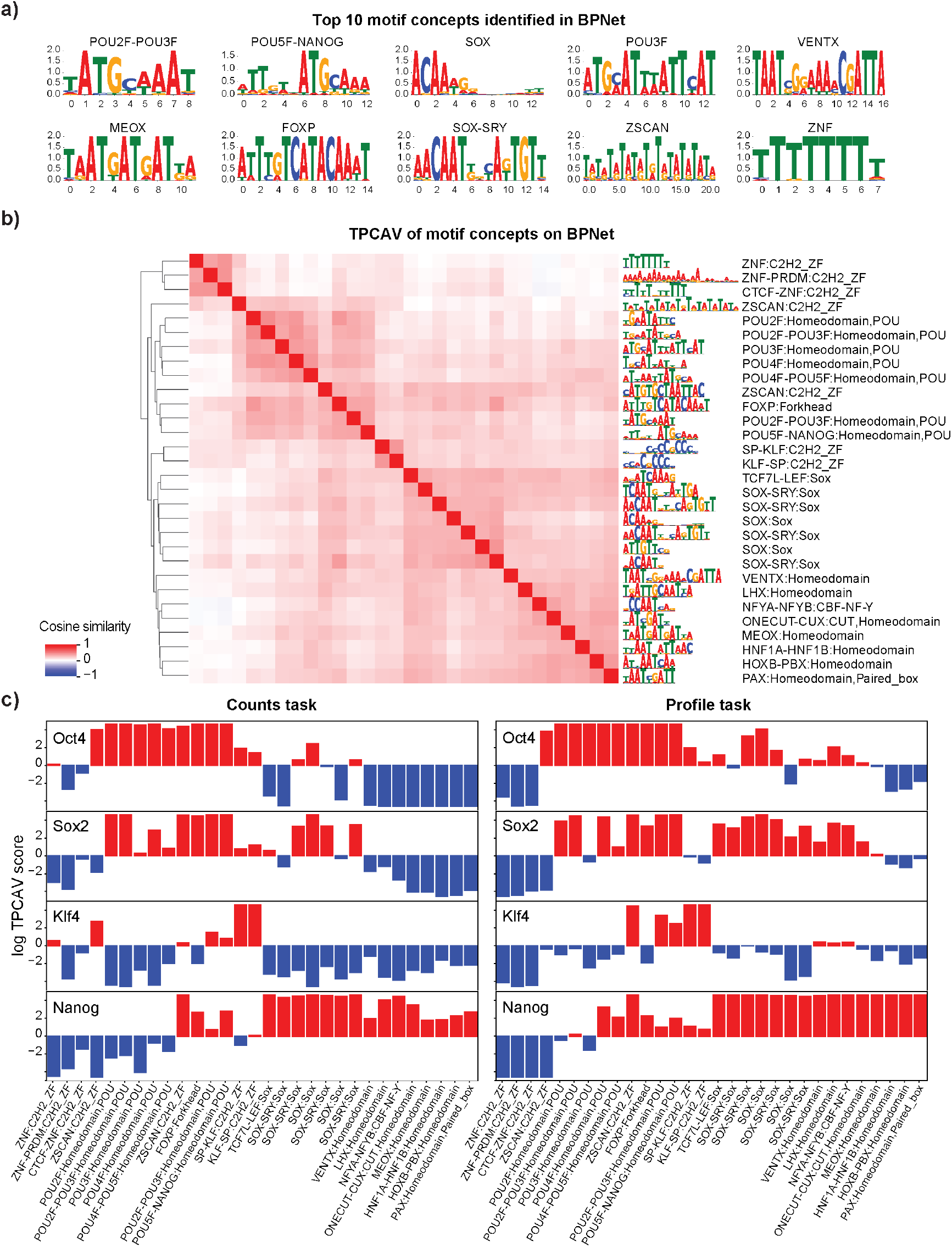
Concept specific attribution map reveals differential usage of concepts across prediction heads in BPNet. **a)** Top 10 motif concepts identified in the last shared convolutional layer by all prediction heads. **b)** CAV cosine similarity heatmap of top 30 motif concepts (F-score > 0.9) in BPNet. **c)** Log TPCAV scores of top 30 motif concepts (F-score > 0.9) across all prediction tasks, bar color indicates whether a concept is positive (red) or negative (blue) influence.

Beyond the expected cognate motif concepts, several homeodomain-related concepts showed specific positive influence in the Nanog count task (**Figure 4a**), consistent with the presence of a homeodomain in the Nanog protein^29^. With respect to repeat concepts, in addition to simple repeat concepts that may reflect GC-content variation or similarity to cognate motif patterns, multiple LTR and SINE element concepts displayed task-specific activity. For example, multiple SINE B1 and SINE B2 elements positively influence Klf4 count predictions specifically. LTR_IAPEz-int and LTR_RLTR17B_Mm show task-specific positive influence for Nanog and Oct4 count tasks, respectively (**Supplementary Figure S7**). The differences between count and profile tasks are primarily observed in low-complexity region and simple repeat concepts. This may indicate that repeat-associated information is more important for accurately shaping the profile distribution, whereas it plays a less critical role in count prediction tasks (**Supplementary Figure S7**).

While the TPCAV score indicates how the model’s predictions change when the input is perturbed toward a given concept direction, thereby revealing the model’s responsiveness to each concept, it does not capture how frequently or strongly each concept is used across TF-bound regions. Given the more complex and heterogeneous embedding space of BPNet, it is particularly interesting to investigate whether the model relies on different concepts amongst peaks of the same TF. The original TCAV study proposed using cosine similarity between input activations and a CAV to rank inputs by their alignment with a concept^11^. However, cosine similarity reflects only the direction of the activations and ignores the magnitude of the attribution score along that direction. As a result, it is not an ideal metric for quantifying the actual usage or contribution of a concept in the model’s predictions. To better understand how each concept contributes to individual prediction tasks, we developed a feature-attribution dissection analysis that isolates the portion of the attribution signal explainable by a specified set of concepts. After obtaining the CAVs for the concepts, we project the PCA-transformed activations onto the given CAV direction and examine the corresponding attribution maps propagated back to the input (**Figure 5a**, see **Methods** for details). This approach enables direct comparison of the usage of each concept across peak regions.

**Figure 5:**
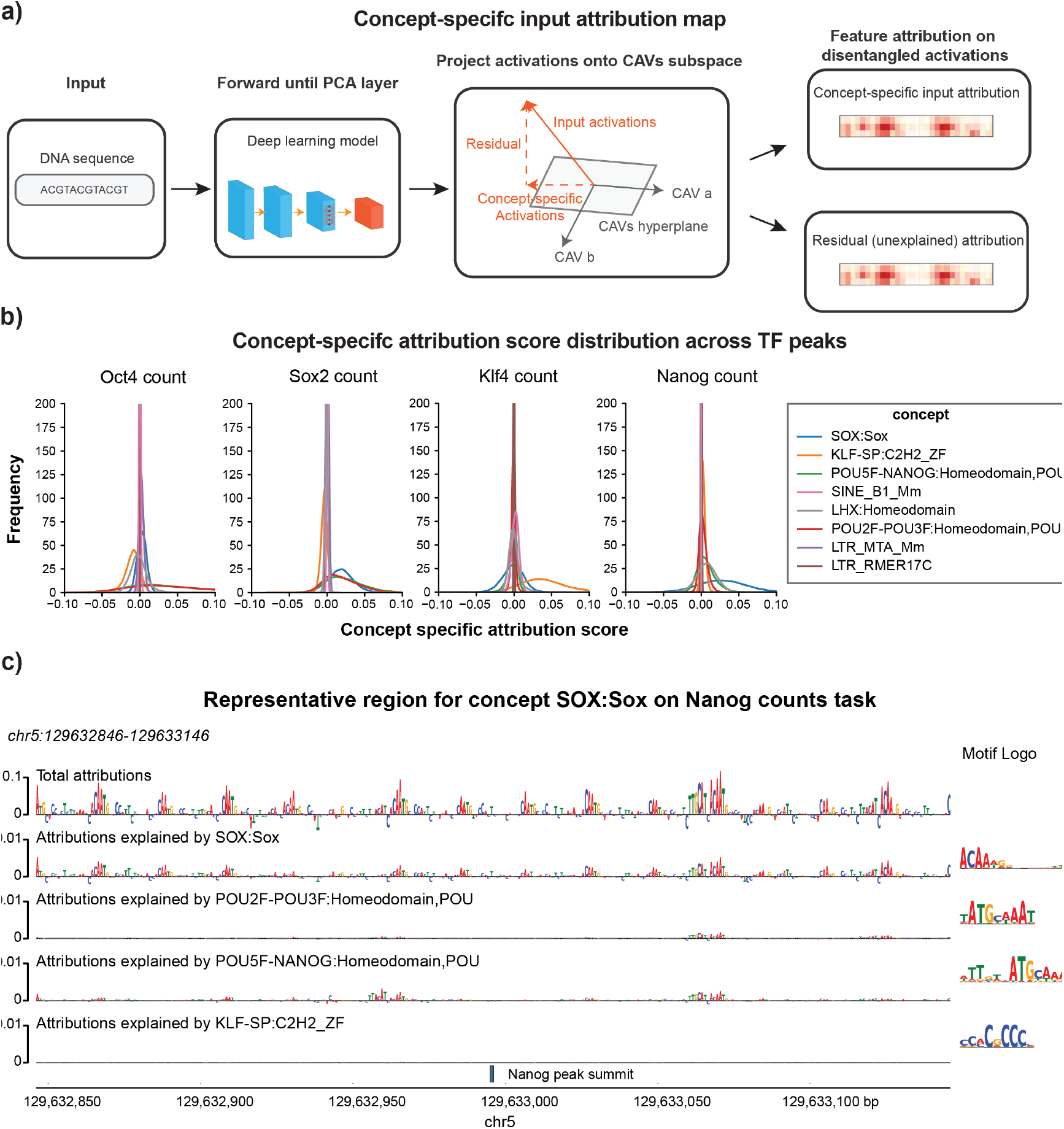
Concept specific attribution map reveals representative peaks of concept in BPNet. **a)** Overview of extracting concept specific input attribution map. **b)** Distributions of the sum of the concept specific input attribution map scores for peaks of each factor, selected concepts show differential TPCAV scores across tasks. **c)** Example of the concept specific attribution map on a top representative region of concept SOX:Sox in Nanog count prediction task.

We selected concept CAVs that displayed differential activity across BPNet’s four count-prediction tasks and examined their contribution patterns. First, we evaluated the distributions of attribution scores projected onto each CAV. These distributions revealed that, for most prediction heads, the cognate motif concept was the most heavily used among peak regions. The notable exception was the Nanog count task, where SOX:Sox contributed the strongest attribution signal, whereas POU5F–NANOG:Homeodomain did not exhibit markedly elevated usage (**Figure 5b**). Interestingly, this observation suggests that the concepts with positive contributions identified in BPNet do not necessarily occur in a large fraction of TF peaks, indicating that concepts with strong influence on model predictions are not always utilized across binding sites.

We next asked whether peaks could be ranked according to their usage of the top cognate motif concept. We ranked each peak set by the sum of the corresponding top cognate motif concept specific attributions for count tasks in each region (**Supplementary Table S4**). Interestingly, the peaks with the highest usage for each concept typically contain multiple instances of the corresponding motif and these motif instances appear in a periodic pattern where each motif copy contributes similar degrees of attribution to the prediction (**Figure 5c** and **Supplementary Figure S8**). This observation is consistent, to some extent, with the original BPNet paper, which reported periodic motif soft syntax among OSKN cognate motifs^1^.

To assess whether the peak rankings capture meaningful biological signal, we selected the top and bottom 3,000 peaks for each factor based on usage of the most influential concept and examined their distances to the nearest TSS. Notably, Klf4 peaks ranked by KLF/SP:C2H2_ZF and Nanog peaks ranked by SOX:Sox showed clear differences between the top and bottom groups: the top-ranked Nanog peaks tended to reside farther from TSSs, whereas the top-ranked Klf4 peaks were, on average, closer to TSSs than their bottom-ranked counterparts (**Supplementary Figure S9**). We further performed GREAT analysis on the ranked peak sets. The results revealed distinct pathway enrichments associated with concept-based peak ranking. Among Nanog peaks, the top 3,000 were enriched for histone methylation and proliferation-related pathways, whereas the bottom 3,000 were more strongly associated with RNA-related processes (**Supplementary Figure S10a**,**b**). For Klf4, chromatin assembly pathways were highly enriched among the top-ranked peaks, while apoptosis-related pathways appeared more frequently among the bottom-ranked peaks (**Supplementary Figure S10c**,**d**).

In summary, TPCAV can effectively find concepts that are associated with each prediction task in multi-headed models, as evidenced by the discovery of informative motif concepts that are consistent with the original BPNet study. By specifically extracting the attribution scores aligned in the selected concept CAV direction, we can extract regions that are representative of the concept.

## Discussion

In this manuscript, we introduce TPCAV, a modified version of the concept-based interpretation method TCAV that is tuned for interpreting genomic deep learning models. To our knowledge, this is the first systematic attempt to analyze deep learning models for genomic tasks using concept-based attribution. Our results show that integrating a PCA transformation into the feature-embedding space substantially improves the stability and reliability of concept-activation measurements. We also demonstrated that TPCAV can be used to assess the influence of multiple types of biological concepts within genomics models, including concepts based on cis-regulatory motifs, genome annotations, and epigenomic signals. Additional biological concepts can be easily defined by collecting a set of DNA sequences or genomic regions that exemplify the concept and another set of sequences that do not contain the concept. We also demonstrated that TPCAV provides comparable DNA motif-level interpretations to TF-MoDISco, although one key difference between methods is that TPCAV interprets the influence of a collection of known motifs whereas TF-MoDISco can discover motifs *de novo* from sequence attribution patterns.

The flexibility of the concept-based attribution framework enables interpretation of biological concepts that are not easily represented by DNA motifs. For example, using DNA repeat concepts, we found that low-complexity GC-rich elements play a prominent predictive role in many single-TF binding models. We also demonstrated that TPCAV can interpret concepts based on non-sequence features in multimodal frameworks such as maxATAC. As expected, TPCAV correctly identified the relevant open-chromatin concepts in these models. Interestingly, GC-rich sequence concepts tended to have negative influence in maxATAC models, in contrast to their positive role in sequence-only models, underscoring how the same biological feature may assume different roles in a predictive task depending on model architecture and training context.

Application to the shared embedding space of BPNet demonstrated that TPCAV can separate the influence of biological concepts on each predictive task in a multi-task model. For example, and intuitively, TPCAV finds that each TF binding prediction task in BPNet is influenced by appropriate cognate motif concepts. To quantify the actual usage of each concept within TF peaks, we developed a feature-attribution dissection analysis that extracts concept-specific attribution maps. Ranking TF peaks by motif concept usage revealed meaningful subclasses of peaks, reflecting internal heterogeneity within TF binding events.

A key advantage of concept-based attribution is its independence from input format and model architecture, making TPCAV highly flexible and generalizable. We demonstrated that TPCAV can be applied to interpret tokenized-input DNA foundation models, including DNABERT-2, the Nucleotide Transformer, and HyenaDNA. To our knowledge, no other global interpretation methods can currently be applied to interpret tokenized-input models. We also demonstrated that TPCAV can be used to benchmark the degree to which pretrained foundation models have learned or encapsulate a diverse set of biological concepts. Applying such a benchmarking approach using our DNA motif concept database across three pretrained foundation models uncovered striking differences in the representation of cis-regulatory motifs in their embedding spaces. DNABERT-2 showed the strongest representation of known TF-binding motifs in its embedding space, whereas larger models such as the Nucleotide Transformer and HyenaDNA performed substantially worse. Parameter scale did not explain these differences, suggesting that training data composition or training strategies may be the primary determinants. It is worth noting that our results measure differences in how foundation models linearly encode motif information in their embedding spaces, which is not necessarily correlated with overall model performance after models are finetuned for downstream tasks.

As genomics models rapidly scale, concept-level interpretation will provide a flexible approach for assessing model reliability, identifying failure modes, and generating experimentally testable hypotheses. The diversity of model architectures and training paradigms makes it challenging to uniformly interpret and compare what different models learn when applied to the same task. The flexibility of TPCAV, which imposes no constraints on input format or model architecture, makes it particularly well suited for this landscape. Moreover, the community has accumulated extensive domain knowledge, and many established concepts (e.g., GO terms, chromatin states) can naturally be incorporated as concept sets. This positions TPCAV as a highly extensible framework for models trained on a wide range of biological tasks.

In addition to post-hoc interpretation, some studies have explored incorporating concept examples directly during training to regularize embedding spaces^30,31^. Such approaches may be especially valuable in biology, as explicitly separating known biological concepts during training could help highlight novel, previously uncharacterized features learned by deep learning models.

In summary, TPCAV provides a powerful complement to current deep learning model interpretation practices in genomics. The research community has established rich knowledge about gene regulatory concepts, and integrating these concepts into model interpretation can help distinguish genuinely novel insights from patterns that are already well understood.

## Methods

### Testing with PCA-transformed Concept Activation Vectors (TPCAV)

TCAV^11^ uses concepts to interpret deep learning models, where each concept is defined using a set of samples that exemplify the concept. Concept examples are presented to a neural network model, along with control samples that do not contain the concept. We train a linear support vector machine (SVM) classifier using stochastic gradient descent (SGD) with L2 regularization at a selected layer *l* to distinguish the concept examples from control samples.

Hyperparameters are tuned via grid search over the regularization strength {10^−2^, 10^−4^, 10^−6^}. If concept examples can be effectively classified from random samples, we can assume that the model embedding space has encapsulated knowledge relating to that concept. Classifier performance is measured here via the F-score, defined as the harmonic mean of precision and recall

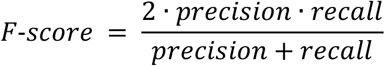

The vector orthogonal to the decision hyperplane of the linear classifier is termed the Concept Activation Vector and represents the direction of the concept in the layer’s embedding space. By taking the dot product of the CAV with the model layer attribution scores for data points of interest, we can assess how the predictions of the model would change when moving towards that concept from the baseline conditioned on the data points of interest (**Figure 1a**).

In practice, deep learning models trained for genomic tasks, such as TF binding prediction, often exhibit redundant parameters, resulting in highly correlated activations at layer *l* that may have either positive or negative effects on the final prediction. For instance, a model trained to predict FOXA1 binding may contain five activations that all represent FOXA1 cognate motifs in the embedding space, but with weights [+10, +8, +3, +1, −5]. Although the negative influence of the 5th activation is compensated by the other four in the final output, a linear classifier trained to capture the FOXA1 concept may mistakenly emphasize this negatively weighted activation simply because all five activations serve as strong discriminative features for the concept. This can lead to aberrant TCAV interpretation of a concept representing the FOXA1 cognate motif.

To address the issue of correlated feature activations, we apply a PCA transformation to the embedding space as an additional operation after layer *l*. For each concept to be evaluated, we sample a small set of examples (typically 10) for each concept and perform a PCA transformation by SVD decomposition on their standardized activations:

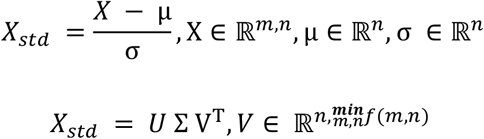

Where *m* is the batch size and *n* is the dimension size after flattening the embedding space. We retain the standardization parameters μ, σ and transformation matrix *V* to transform future samples into the PCA projected space. In case that we could not obtain the full rank transformation (*m* < *n*), the remaining residual not explained by PCA is also kept and concatenated to the PCA projected vector

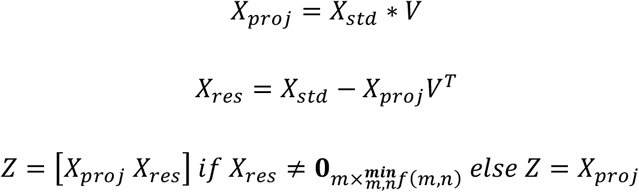

The concept linear classifier is then trained on the final concatenated PCA transformed feature vector and residual vector *Z* between concept examples and control examples. The training is accomplished by SGDClassifier implemented in scikit-learn^32^, the learning rate is set as ‘optimal’ and we performed a grid search on alpha parameters ranging in [1*e* − 2, 1*e* − 4, 1*e* − 6] for every concept

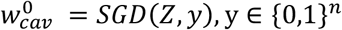

#### Layer attribution

Layer DeepLift attribution^9^ was performed using Captum^33^. For each test region, we sampled 10 random genomic regions as baseline samples. We obtained the layer DeepLiftSHAP score by averaging the LayerDeepLift scores computed against all 10 random background regions. Usually, attribution scores are computed by multiplying multiplier/gradient with the difference between input and baseline^34^:

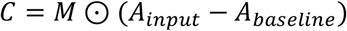

Here, the difference between the input and the baseline serves two key purposes: (i) it reflects the actual magnitude of the attribution signal, highlighting what drives the change in prediction between the input and the baseline; and (ii) it preserves the sign of this difference, capturing whether the contribution is positive or negative. However, in TPCAV, the sign of the final TPCAV score is determined by the decision boundary of the classifier on concept examples and control examples, thus we only consider absolute magnitude of the difference between inputs and baselines in the final TPCAV score calculation

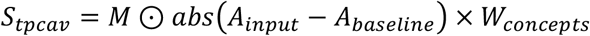

#### Concept-specific attribution maps

To obtain the attribution map specifically explained by given CAVs during the forward process of the model, we injected an additional operation to project the PCA-transformed activations onto the column space of the given CAVs:

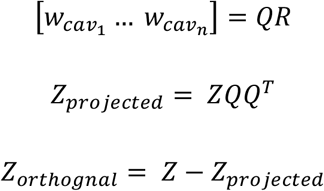

Here, *Z*_*projected*_ denotes the projection of the activations onto the column space of the given CAVs, while *Z*_*orthognal*_ represents the residual orthogonal component. During DeepLIFT attribution, we explicitly set the multipliers associated with *Z*_*orthognal*_ to zero so that only the multipliers linked to *Z*_*parallel*_ are propagated back to the input

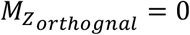

In this way, the resulting input attribution map reflects only the contributions that can be explained by the column space of the provided CAVs.

### Motif concept sensitivity score calculation

To compute the motif concept sensitivity score, we first calculate F-scores for each motif concept across varying numbers of motif insertions. We then compute the area under the curve (AUC) of the F-scores with respect to the number of motif insertions. To account for bias introduced by motif GC content, we compute the weighted GC information content *IC*^*GC*^ as:

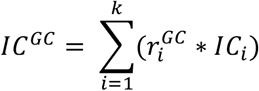

Where 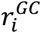 denotes the GC ratio at position *i, IC*_*i*_ is the information content as position *i*, and *k* is the motif length.

We then fit a linear regression model between *IC*^*GC*^ and the AUC of F-scores to remove the variance explained by GC content. The residual from this regression is defined as the motif concept sensitivity score.

### Biological concept database construction

#### Motif concepts

We used the non-redundant human motif clustering database from Jeff Viestra’s lab^13^ to ensure comprehensive coverage of candidate motifs. To construct concepts representing motif instances, we first randomly sampled 5000 genomic regions. For each motif concept, we randomly inserted multiple motif instances, generated from the corresponding position weight matrix, into each region, ensuring no overlaps between inserted motifs. Motifs were sampled in either the forward or reverse orientation.

For single TF prediction models trained on ENCODE datasets and maxATAC^7^, we inserted 12, 24, or 36 motifs into a 1024bp input window. For BPNet^1^, we inserted 12, 24, and 36 motifs into a 1000bp input window. For DNABERT-2^19^, Nucleotide Transformer^23^, and HyenaDNA^24^, we inserted 3, 6, 12 motifs into a 512bp input window.

#### Open chromatin concepts

We constructed open chromatin concepts by randomly sampling 5,000 DNase-seq or ATAC-seq peaks for each cell line, using datasets downloaded from ENCODE^35^. Each region’s DNase-seq or ATAC-seq profile was paired with sequences of a random genomic region to separate the influence of the epigenomic signal from the DNA sequence features corresponding to that signal.

#### DNA repeat concepts

Repeat concepts were derived from the RepeatMasker BED annotations^15^. For each repeat class defined by RepeatMasker, we sampled 5,000 instances as concept regions and excluded repeat types with fewer than 5,000 available examples.

### Single TF binding prediction model dataset construction and training

#### Model architecture

The model architecture is inspired by Enformer^3^ and implemented using Pytorch Lightning framework^36,37^, It consists of four parts (**Figure 1c**, model architecture diagram generated by NN-SVG^38^):

1. The first part of the model is a single 1-D convolutional layer that includes 128 filters of size 25bp.
2. Then there is a convolutional tower composed of four design blocks, each block has a 1-D convolutional layer of *n* filters of kernel size 5 and skip connection, a 1-D convolution layer of 2*n* filters of kernel size 1, and a 1-D Max Pooling layer of kernel size 2. n is a function of the index *i* of the block in the convolutional tower

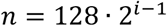
3. After the convolutional tower, a positional encoder is applied to the activations, followed by a transformer encoder layer of 8 attention heads and 1024 feedforward dimensions.
4. Finally, a 1-D convolution layer reduces the channel size to 1, then a fully connected linear layer predicts the final logit.

The full list of operators is available as listed in the following table:

| Operator | Params | Function |
| --- | --- | --- |
| Conv1d: 128 filters; kernel size 25; stride 1 | 1 layer | Extract short local sequence patterns |
| Conv1d: $n$ filters; kernel size 5; stride 1<br>Conv1d: $2n$ filters; kernel size 1; stride 1<br>MaxPooling: kernel size 24 | 4 layers<br>$n = 128 \cdot 2^{i-1}$ | Convolutional tower for capturing local features |
| Transformer encoder layer: 8 attention heads; 1024 feedforward dimensions | 1 layer | Capture long-range interactions |
| Conv1d: 1 filter | 1 layer | Dimension reduction |
| Linear: 1 output feature | 1 layer | Final prediction |

Each convolutional layer or linear layer is followed by LeakyReLu activation and 1-D BatchNormalization layer, except the last linear layer.

#### Dataset

Positive samples were generated by randomly shifting the 256-bp window around ChIP-seq peak summits, ensuring that the model does not overfit to the peak center. Each peak was shifted multiple times to augment the positive training set. Negative samples were 256-bp genomic regions constructed from three sources: (i) purely random genomic regions lacking TF binding, (ii) random unbound regions matched to the GC content distribution of peak regions, and (iii) flanking regions surrounding peaks. Each negative category was sampled at approximately equal size. The test set additionally included a separate collection of random genomic regions to assess genome-wide performance. Finally, all sample regions are expanded to 1024-bp long input windows with their labels determined by the center 256-bp region.

#### Training, validation, and testing

During training, each batch was balanced to contain equal numbers of positive and negative examples to prevent the model from biasing toward the more prevalent negative class. For validation and testing, the full sets of samples were used without rebalancing. Model parameters were optimized using the AdamW^39^ optimizer with a base learning rate of 1*e*^−5^. A OneCycleLR^40^ scheduler was applied with a maximum learning rate 1*e*^−4^ of over 40 training epochs.

## Code Availability

TPCAV is available as a python package hosted at Github (https://github.com/seqcode/TPCAV).

## Acknowledgements

This work was supported by the National Institutes of Health grant R35GM144135 (to S.M.) and the National Science Foundation DBI CAREER 2045500 (to S.M.). Any opinions, findings, and conclusions or recommendations expressed in this material are those of the authors and do not necessarily reflect the views of the National Science Foundation. J.Y. was partially supported by Rising Researcher award ICDS_RR25_027636 from Penn State’s Institute for Computational & Data Sciences (RRID:SCR_025154). The authors thank the members of the Center for Eukaryotic Gene Regulation at Penn State for helpful feedback and discussions.

## Notes

### Competing Interest Statement

The authors have declared no competing interest.

### Summary of Updates

Supplementary Tables were missing from the previous submission. Submitting the Supplementary Tables in this revision.

