## Supplementary Figures for "TPCAV: Interpreting deep learning genomics models via concept attribution"

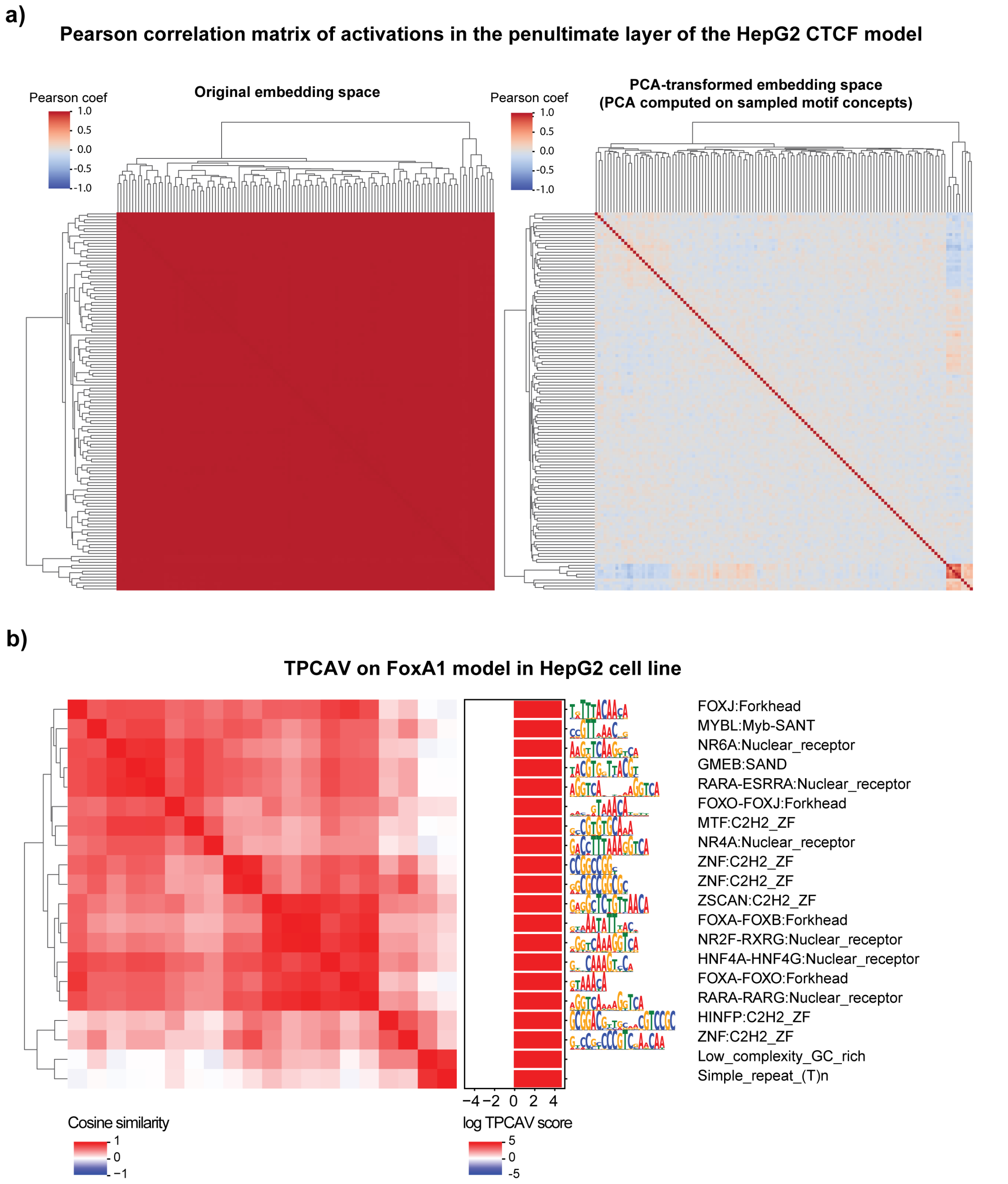


**Figure S1: a)** Pearson correlation of neuron activations before the final dense layer in CTCF peaks and random regions before (left) and after (right) PCA transformation. **b)** CAV cosine similarity heatmap and TPCAV scores of top motif concepts (among top 30 and F-score > 0.9) and repeats concept (F-score > 0.9) in FOXA1 prediction model trained in HepG2 cell line.

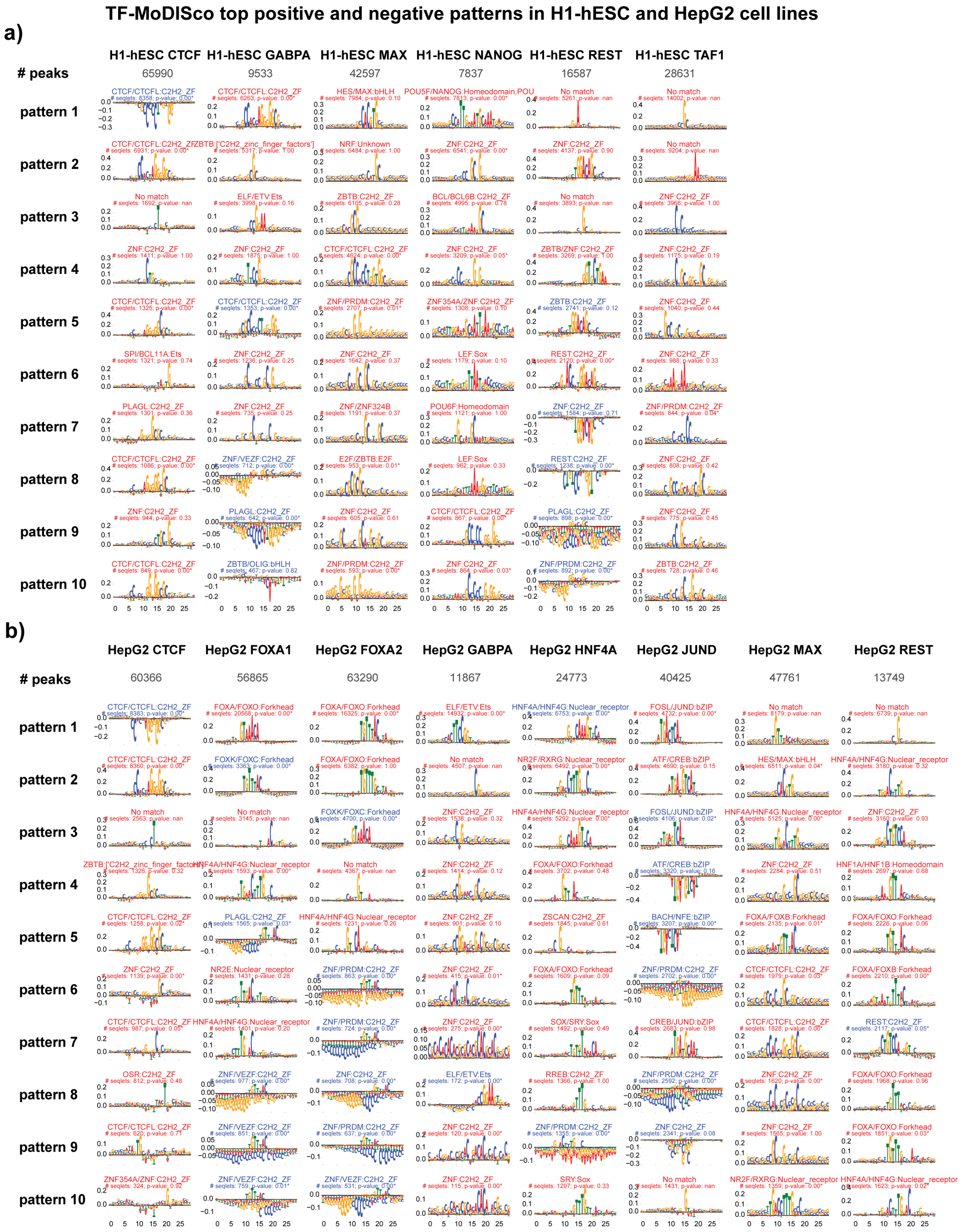


**Figure S2: a)** Top 10 TF-MoDISco sequence patterns in models trained in H1-hESC cells. **b)** Top 10 TF-MoDISco sequence patterns in models trained in HepG2 cells. Motif names in red denote motifs found in positive seqlets, motif names in blue denote motifs found in negative seqlets.

**Performance of linear classifiers for the top 30 motif concepts across input window subregions**

Four motif instances inserted into 128-bp bins with a 32-bp stride

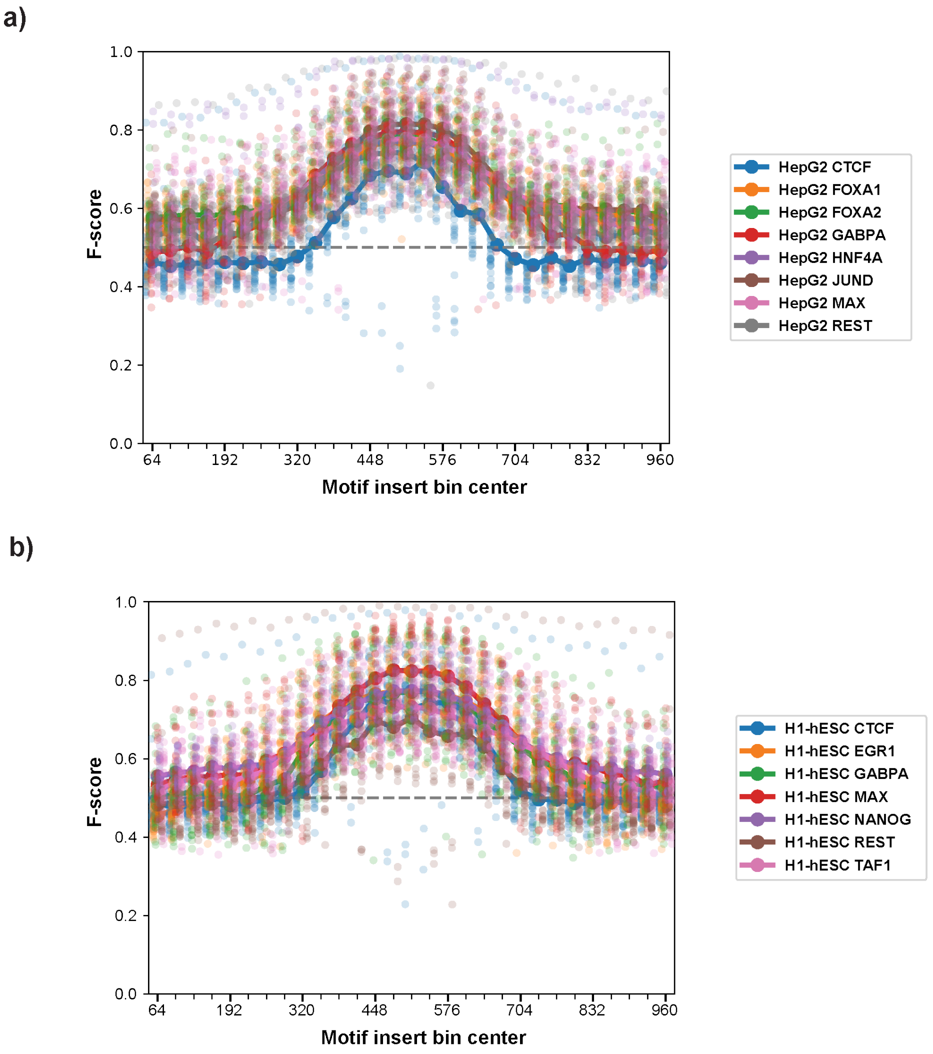


**Figure S3:** Distributions of F-scores of the top 30 motif concepts inserted at varying 128bp subregions inside the input window for models trained in **a)** HepG2 **b)** H1-hESC.

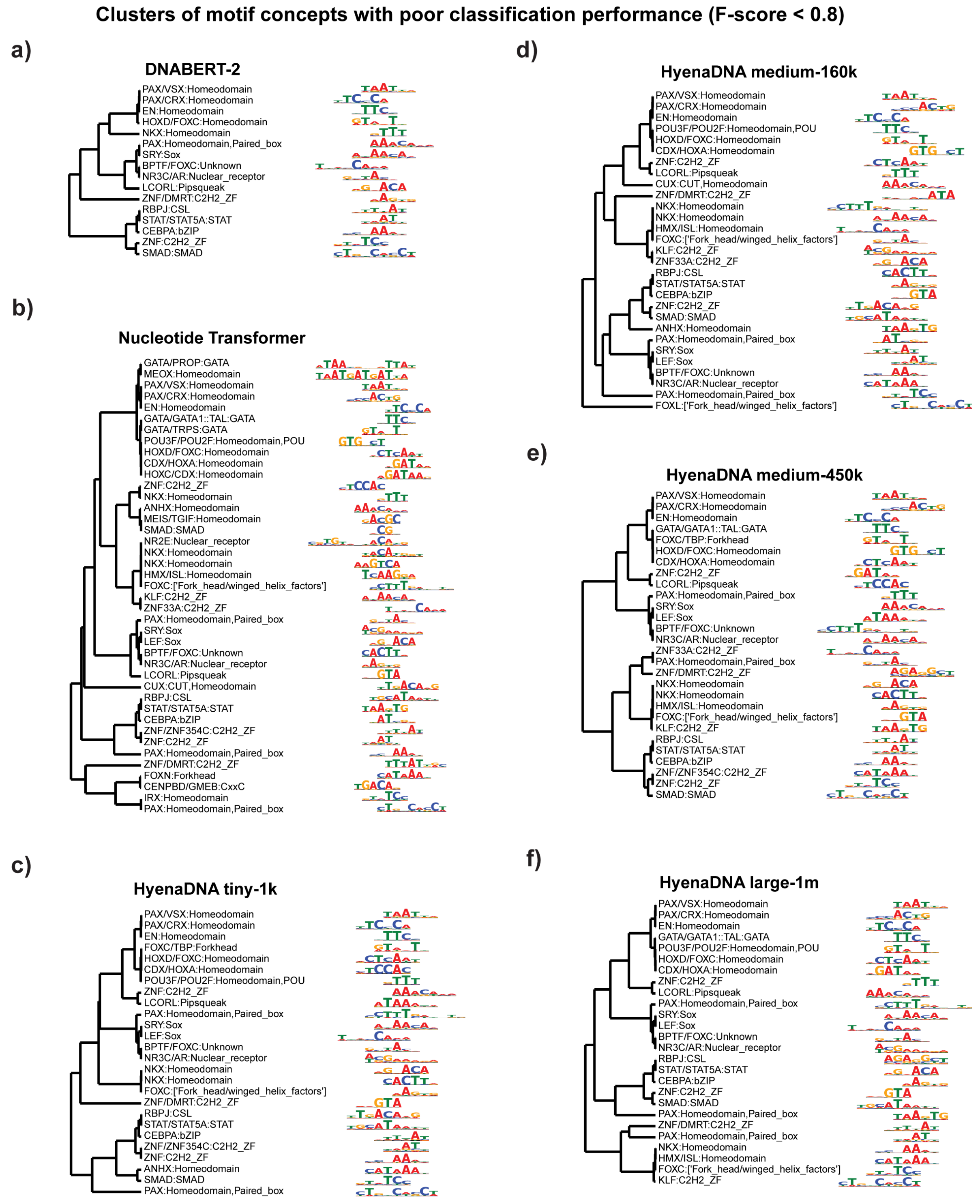


**Figure S4:** STAMP motif clustering and alignment on motif concepts underrepresented (F-score < 0.8 at 12 insertions) in the embedding space of each pre-trained foundation model.

### Top 30 motif concepts on foundation model embeddings

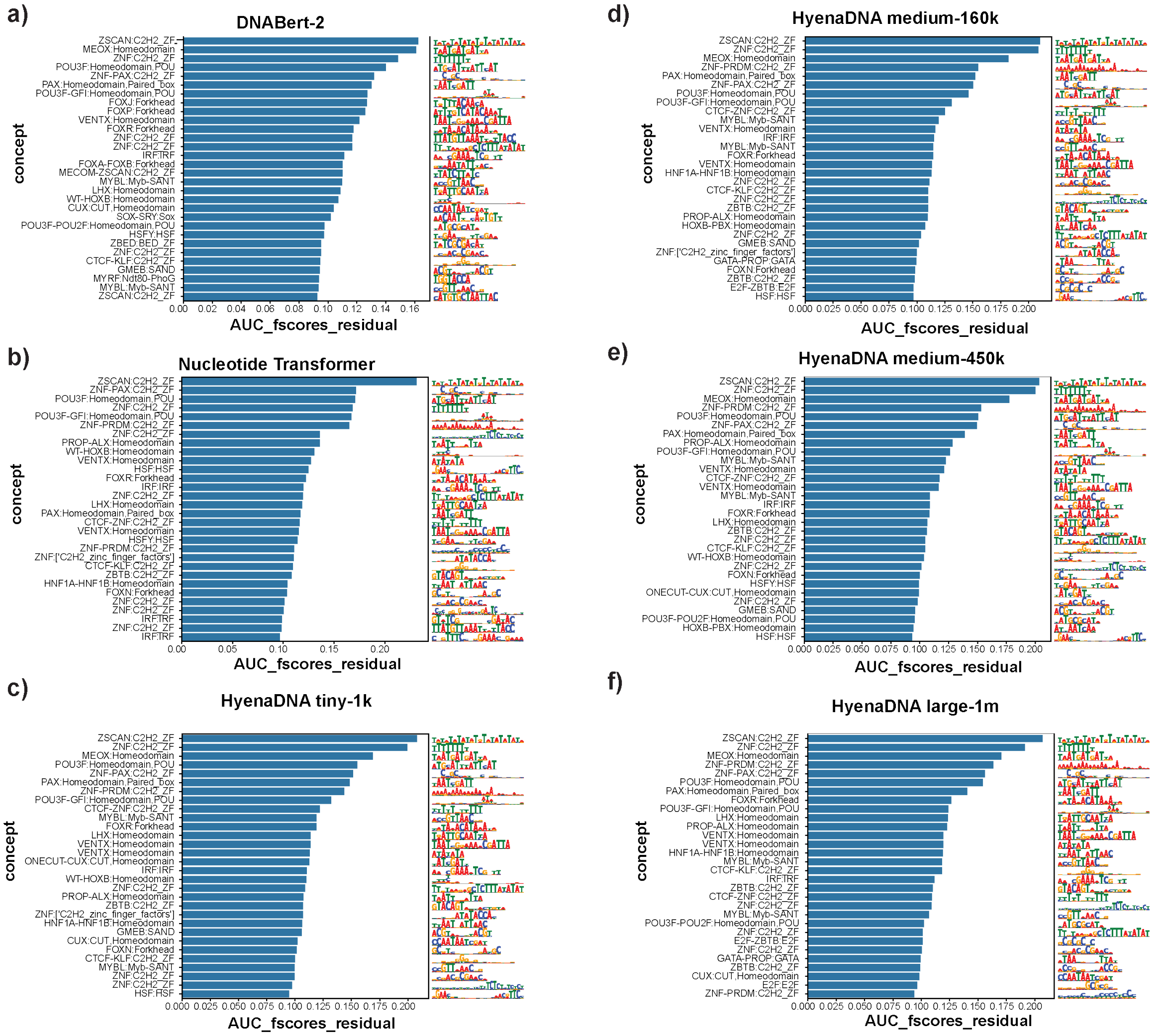


**Figure S5:** Top 30 motif concepts recognized by TPCAV in the embedding space of each pre-trained foundation models, ranked by the residuals of AUC F-scores.

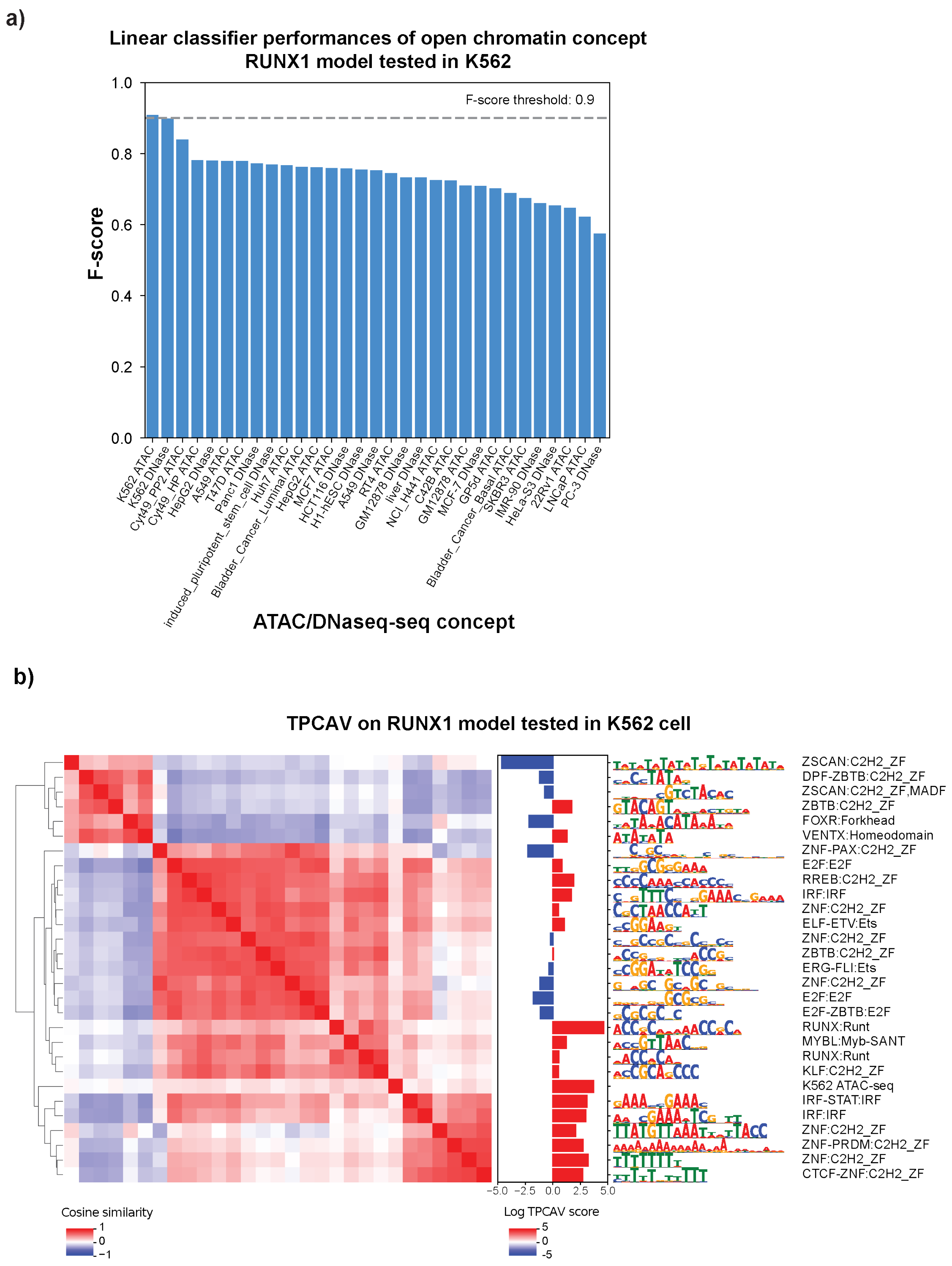


**Figure S6: a)** F-score distribution of open chromatin concepts in maxATAC K562 RUNX1 model. **b)** Cosine similarity matrix between motif concepts (among top 30 and F-score > 0.9) and open chromatin concepts (F-score > 0.90) in maxATAC K562 RUNX1 model.

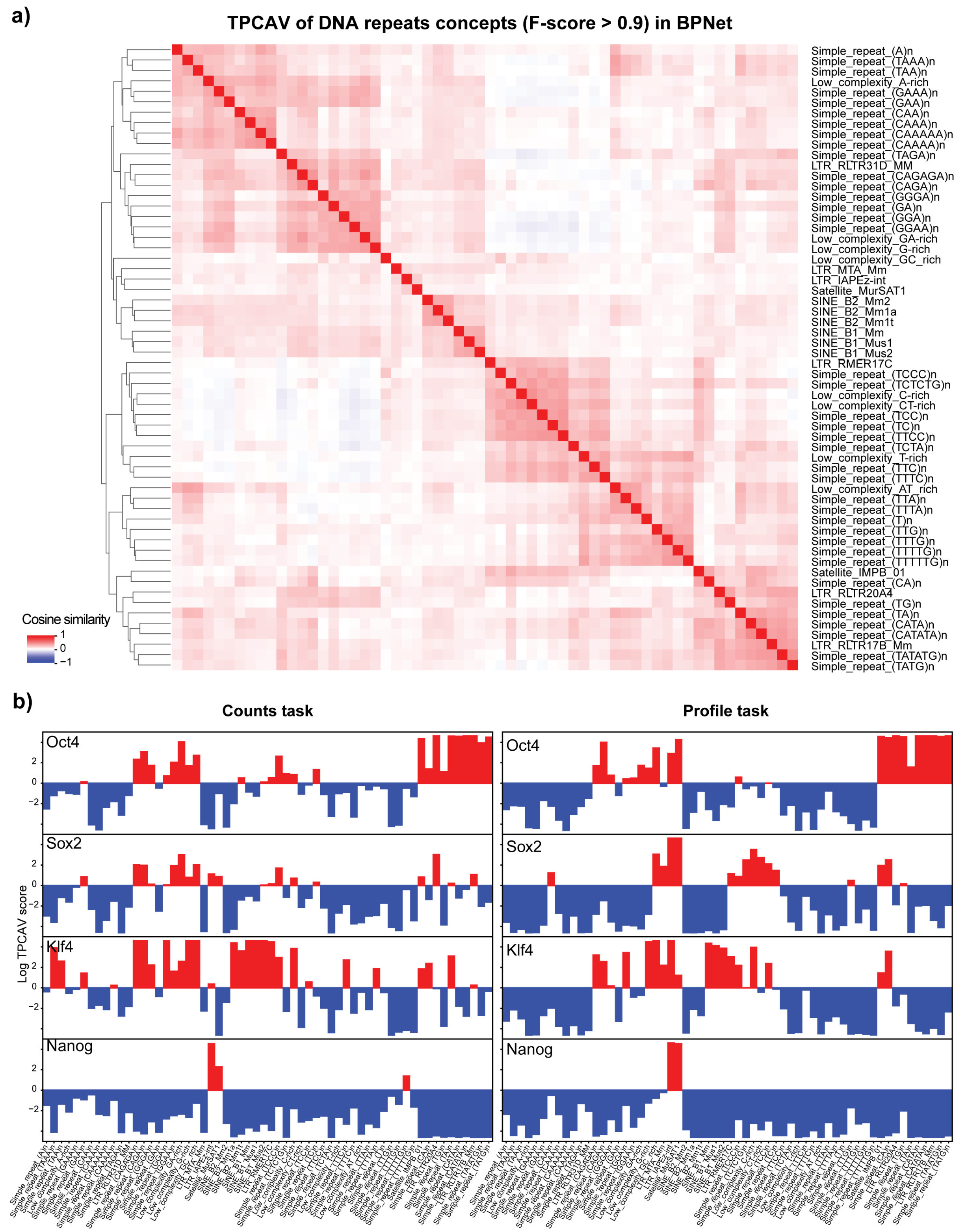


**Figure S7: a)** CAV cosine similarity heatmap of repeat concepts (F-score > 0.90) in BPNet. **b)** TPCAV log ratios of repeat concepts across each prediction head.

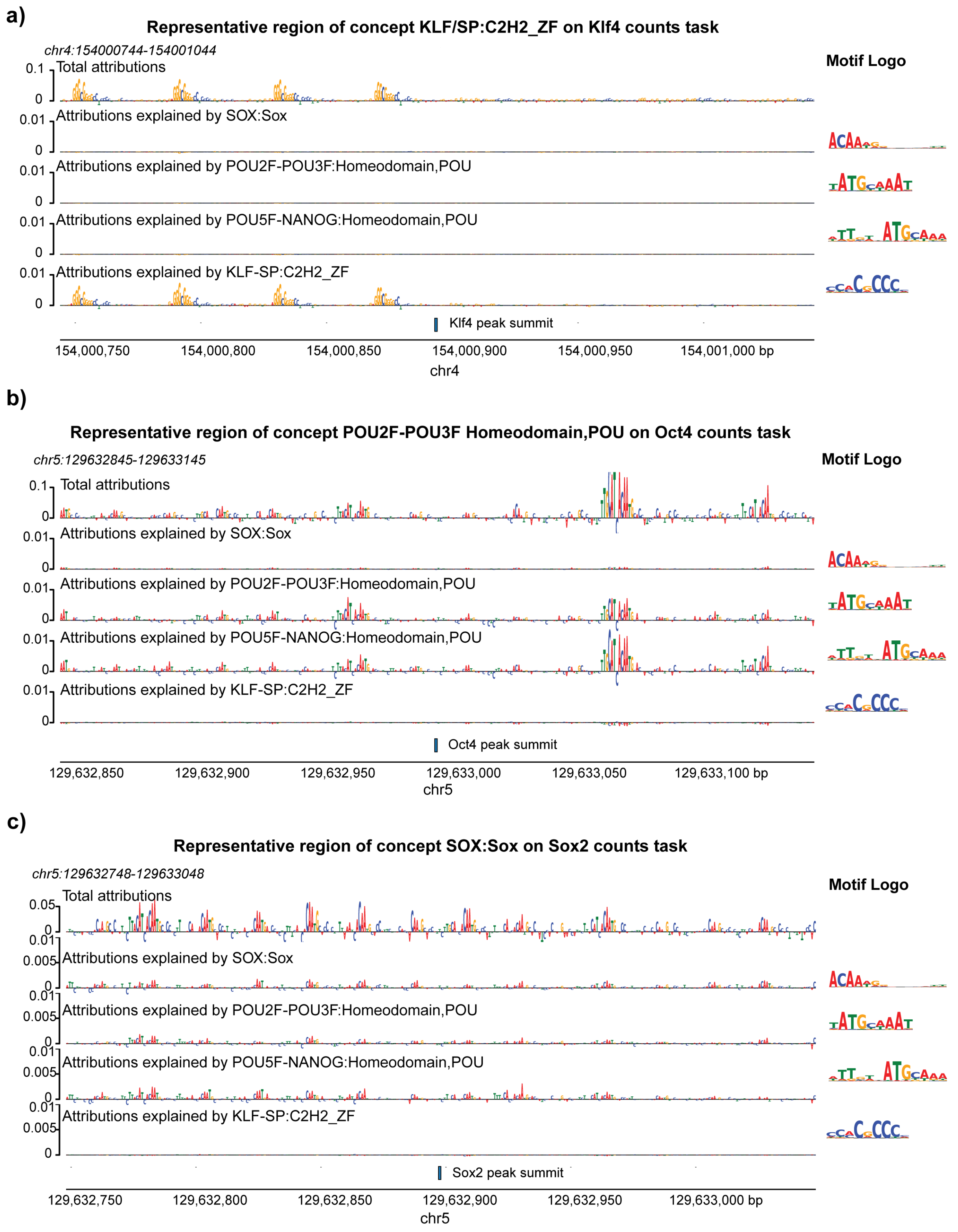


**Figure S8:** Example concept-specific attribution maps for BPNet counts tasks and cognate motifs in top representative regions. **a)** Klf4 counts task and the KLF/SP motif; **b)** Oct4 counts task and the POU2F-POU3F motif; **c)** Sox2 counts task and SOX motif.

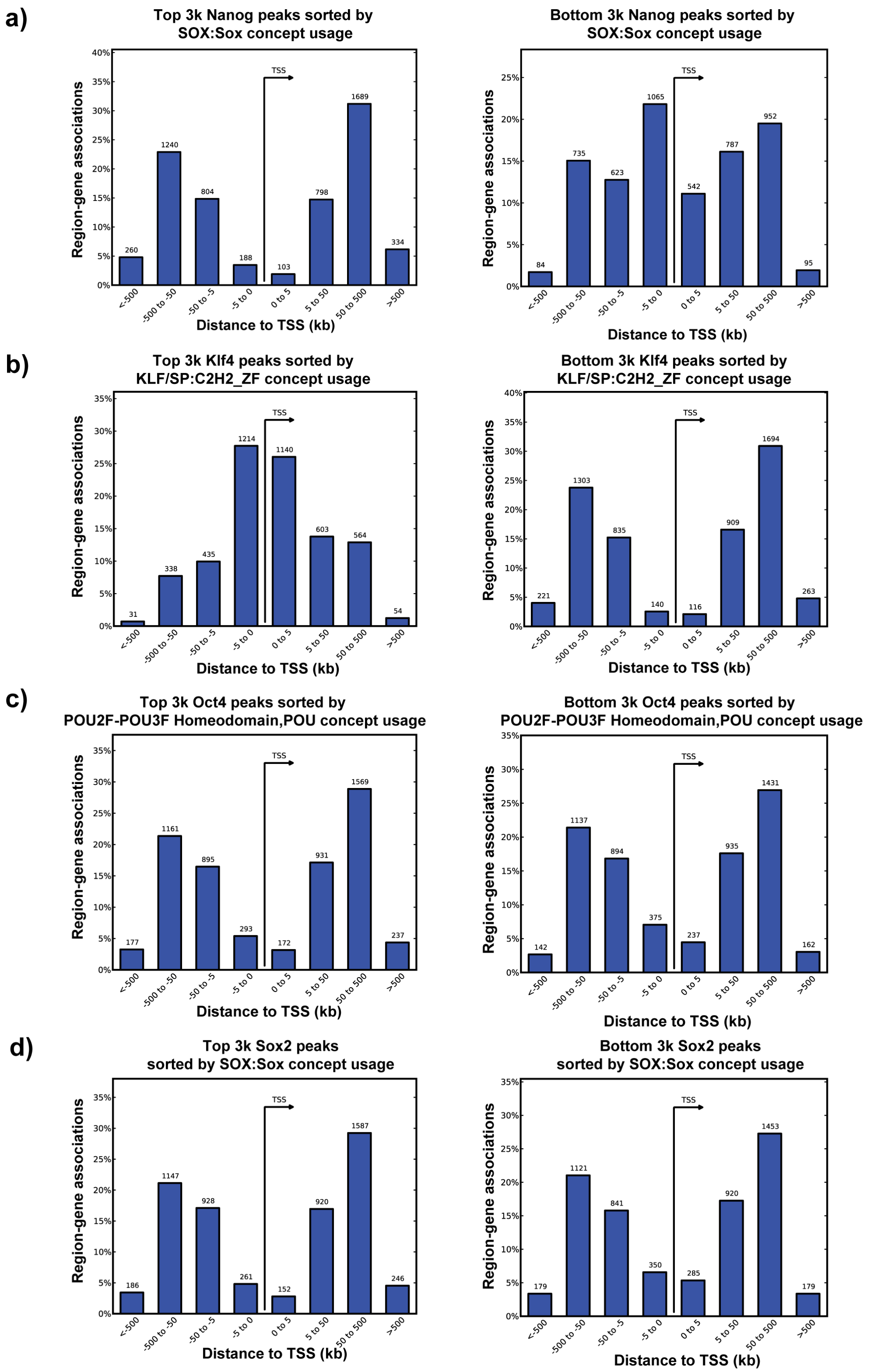


**Figure S9:** Distributions of the distances to the nearest TSSs for top 3k and bottom 3k peaks, ranked by the cognate motif concepts in each peak set.

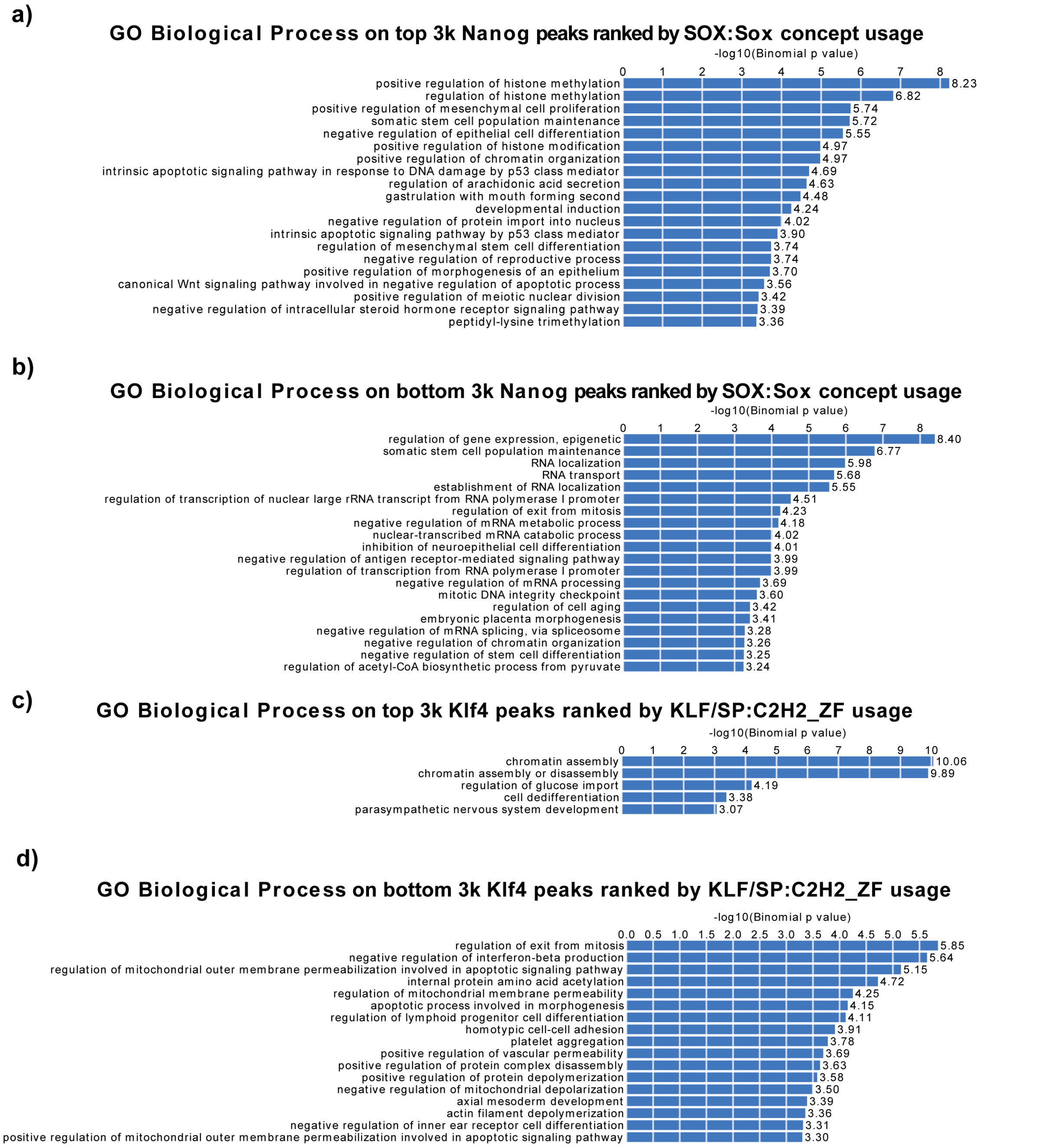


**Figure S10:** Enriched GO:BP pathways on top 3k and bottom 3k peaks, ranked by the cognate motif concepts in Nanog and Klf4 peak set.
